# Prevalence of dual-donating amines in key regions of functional RNAs

**DOI:** 10.1101/2025.06.05.658080

**Authors:** Andrew J. Veenis, Md. Sharear Saon, Philip C. Bevilacqua

## Abstract

RNA performs many critical functions nearly all of which are enabled by complex hydrogen bonded structures. Nucleotides possess far fewer hydrogen bond donors than acceptors, and only the exocyclic amine can donate two H-bonds, suggesting a specialized role. To assess the prevalence and structural contexts of dual-donating amines within structured RNAs, we created a computational workflow that mines and analyzes experimental RNA-containing structures. We evaluated H-bonding in over 250,000 amines from more than 1,800 structures. Dual-donating amines were found most frequently in G’s where they regularly interacted with diverse pairs of acceptors. In contrast, the dual-donating amines of A’s and C’s were less frequent and they interacted with a more select set of acceptors. For all three nucleobases, amines that were dual- donating had both reduced solvent accessibility and higher atom density relative to amines that were non-donating, indicating a tendency of dual donors to be more buried and help compact the RNA. Moreover, analysis of RNA pseudo-torsion angles revealed that dual-donating amines are enriched in two non A-form conformations, both of which are present in S-motifs found in the sarcin-ricin loop of rRNA. We find that dual-donating amines populate additional structural motifs including the GNRA tetraloop receptor, the kink-turn, and the WC/H A-minor motif, which are present in the self-splicing group I intron, the SAM riboswitch, and the poly(A)-bound ENE. We suggest that dual-donating amines may enhance interactions by reducing conformational entropy loss as well as strengthening nearby H-bonds.

## Introduction

RNA serves myriad functions in biology, including in gene regulation, splicing, tRNA processing, and translation and does so by adopting complex secondary and tertiary structures (Leamy et al. 2016). These structures often contain single stranded segments enriched in bulges, loops, and junctions that can interact with one another by an array of interactions including hydrogen bonding, pi-stacking, and metal coordination. Such interactions can compact the RNA, leading to function.

One method of assessing interactions across varied RNA-containing structures is cheminformatics, which entails computationally mining and analyzing molecular data. The Protein Data Bank (PDB; rcsb.org) contains a wealth of data on structured RNAs, with 8,595 RNA-containing entries as of April 10, 2025 (Berman et al. 2000). Over the last decade, the number of high-resolution RNA structures has increased dramatically, a trend that should continue given the ever-increasing interest in RNA and the ongoing improvement in structure determination methodologies (Ma et al. 2022). Using cheminformatic approaches, these structures can be surveyed using Python along with a host of computational tools such as PyMOL, Biopython, and GEMMI (Hamelryck and Manderick 2003; Cock et al. 2009; Wojdyr 2022). Indeed, our group has contributed to cheminformatics of RNA in multiple ways, including identifying the prevalence of *syn* bases in the active sites of RNAs (Sokoloski et al. 2011), determining common catalytic strategies of small ribozymes (Seith et al. 2018), and reporting alternative protonation in the formation of RNA structures (Saon et al. 2025). Knowledge of the underlying RNA chemistry, such as bond distances, angles, molecular geometries, and intermolecular interactions, drives the design of computational workflows to analyze the structural data.

Combining information extracted from hundreds of structures, leading to hundreds of thousands of interactions, enables a comprehensive review of RNA interactions and a better understanding of how they might promote RNA form and function.

Herein, we applied a cheminformatic approach to dual-donating amines— exocyclic amines engaged in two H-bonds (see full definition in the Methods)—within structured RNAs. Our analysis includes evaluations of 1) nucleobase types of dual- donating amines and their corresponding H-bond acceptors, 2) dual-donating amine burial within their encompassing structures, and 3) correlations between amine dual- donation and select RNA conformations. We consider more than 1,800 RNA-containing PDB entries enumerated in the 3.386 release of the actively maintained Representative Sets of RNA 3D Structures, which reduces redundancy relative to all available PDB entries (Leontis and Zirbel 2012). Because many structures lack hydrogens, our workflow adds these to the exocyclic amines and then identifies H-bonds on the basis of hydrogen-acceptor distance and donor-hydrogen-acceptor angle criteria. The amines are then sorted into “non”, “single”, and “dual” categories. When combined with other details curated from the structural data, the resulting insights deepen our understanding of dual- donating amine prevalence and context.

## Results

RNA possesses many more hydrogen bond acceptors than donors (Figure 1). Among the four canonical RNA nucleotides, there are 62 lone pairs from oxygen and nitrogen atoms, which can each accept an H-bond, while there are only 12 polar hydrogens that can donate one. Thus, acceptors outnumber donors 5-to-1. Notably, all 28 oxygen atoms in RNA have two lone pairs and so can accept multiple H-bonds; in contrast, only the 3 exocyclic amines, present in A, C, and G, can *donate* multiple H-bonds, making them especially important.

**Figure 1.**
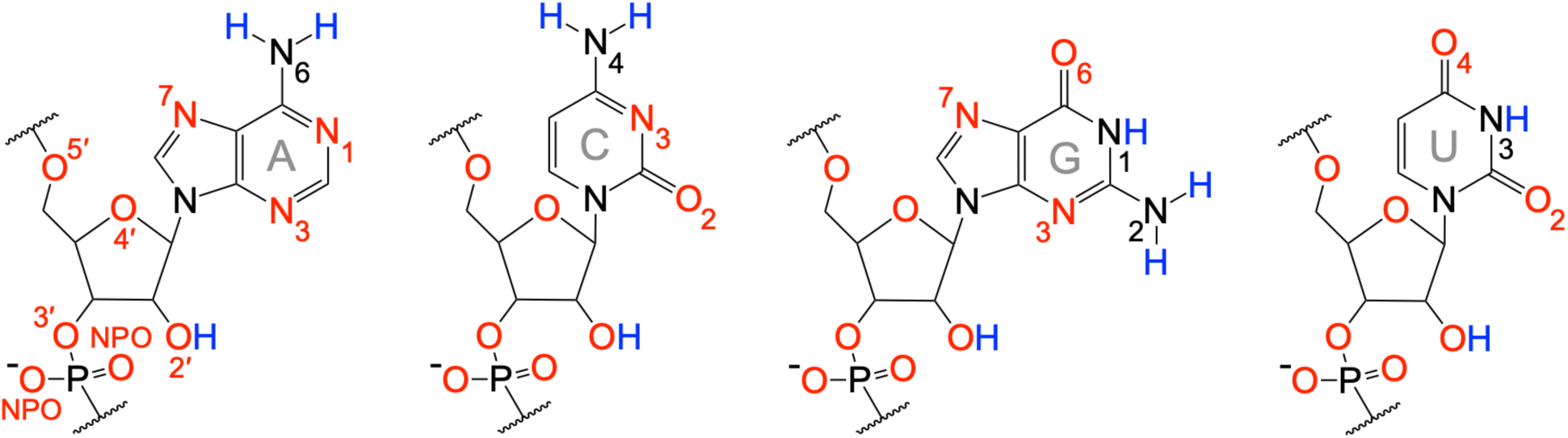
H-bond acceptors vastly exceed donors in RNA. Acceptors and donors are depicted in red and blue, respectively, consistent with partial negative and positive charges. Oxygens can accept two H-bonds via their two lone pairs while deprotonated nitrogens can accept a single H-bond via their single lone pair. The exocyclic amine, which cannot accept an H-bond due to donation of their lone pair into the ring system, can donate two H-bonds via its two polar hydrogens. In contrast, all other nitrogen and oxygen donors can donate only a single H-bond. This counting of H-bonds does not take multifurcated interactions into account (e.g., a bifurcated H-bond), which are enumerated in Supplemental Figure S9. Labels to the key atoms are provided, with the phosphate-sugar backbone key atoms labeled for A only. Only A, C, and G have the dual-donating amine functionality.

### Amine-Acceptor Geometries Inform H-Bond Selection Criteria

We began studying dual-donating interactions with a systematic cheminformatics approach. The 3.386 release of the Representative Sets of RNA 3D Structures, hereafter referred to as the “Representative Dataset”, lists RNA-containing structures that can be downloaded from the PDB and that afford over 250,000 amines to study. Our algorithm queried these amines and determined whether an H-bond forms between each amine and nearby acceptors, independent of the amine being single- or dual-donating. This was accomplished by comparing the measured H-bond distance and angle to user-defined criteria informed by a heatmap of amine-acceptor H-bonding geometries depicted in Figure 2A. In this heatmap, the two key geometries considered are as follows: 1) distance between the acceptor and an amine’s hydrogen (added using PyMOL) and 2) angle between the nitrogen, an amine hydrogen, and the acceptor. To assess these criteria, acceptors near each amine were queried, and their distances and angles were plotted as a heatmap (Figure 2A). Because they would overwhelm the plot due to their presence in canonical base pairing, A(N6)-U(O4), C(N4)-G(O6), G(N2)-C(O2), and G(N2)-C(N3) pairs were excluded from this analysis, although they are included in all subsequent analyses. The resulting heatmap revealed a population towards the middle-right region colored in yellow and green (Figure 2A). The edges of this population guided the selection of the H-bond distance and angle criteria of ≤ 2.5 Å and ≥ 140°, which is represented by the dashed red box, consistent with what would be expected for a typical H-bond (Steiner 2002). Many of the amine-acceptor pairs in the populations outside of the dashed red box likely correspond to weak interactions such as where amine hydrogens are positioned near acceptors due to other, stronger interactions; for example, in the noncanonical WCF GA base pair present in PDB ID 7UW1 (Morgan et al. 2023), the G(N2)-A(N1) pair exhibits a geometry near 2.8 Å and 132°, which falls outside the selected H-bond criteria (Figure 2B).

**Figure 2.**
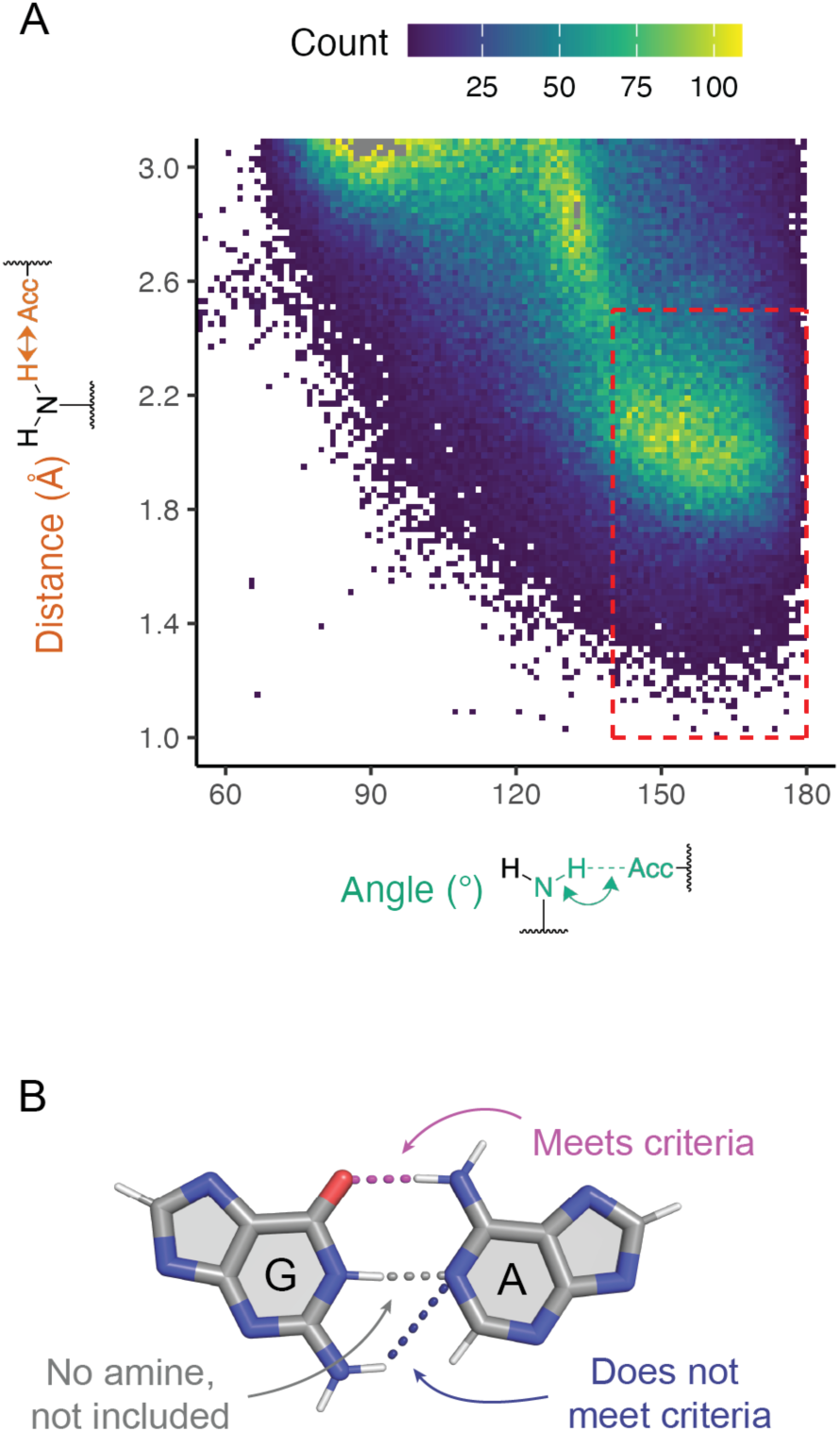
Amine-acceptor pair measurements to determine H-bonding criteria. (*A*) Heatmap depicting amine-acceptor geometries of all amines. Because of their dominance from canonical base pairs, A(N6)-U(O4), C(N4)-G(O6), G(N2)-C(O2), and G(N2)-C(N3) pairs are not included. Any pairs with distance and angle measurements that fall within the dashed red box are considered H-bonding. The points colored grey have counts beyond the maximum of the color scale. (*B*) Examples that meet and do not meet criteria. Depiction of the GA base pair from chain a in PDB ID 7UW1 involving G81 and A76. Magenta: interaction involving an amine that falls within the dashed red box and meets the H-bonding criteria. Blue: interaction involving an amine that falls outside the dashed red box and thus does not meet the H-bonding criteria. Grey: an interaction that does not involve an amine and thus is not included.

### Preferences for Nucleobase Amine Hydrogen Bonding

A first step for understanding the role of dual-donating amines in RNA structure is evaluating the frequency at which A, C, and G amines dual donate. We assessed the structural data on hydrogen bonding of the more than 250,000 amines in the Representative Dataset, with more than 75,000 amines for each nucleobase (Figure 3A). We note that, in contrast to Figure 2, no amine-acceptor measurements that meet the H-bond criteria were excluded. Overall, residues bearing a dual-donating amine are relatively rare, making up only 8.1% (2.1% A + 1.4% C + 4.6% G) of the amines across our dataset. Nonetheless, there are still many dual-donating amines in the Representative Dataset, with 20,478 instances (5,272 A’s + 3,508 C’s + 11,698 G’s). Notably, G’s are highly represented, containing more dual-donating amines than A’s and C’s combined. This enrichment holds even when considering the relatively higher number of G’s in the Representative Dataset (96,177 G’s versus 81,440 A’s and 75,877 C’s), leading to 1 in 8 G’s, 1 in 15 A’s and 1 in 22 C’s being dual-donating. The amine of G resides in the minor groove, while the amines of A and C are in the major groove. Opposite to B-form DNA, the minor groove of A-form RNA is wide and shallow, affording greater accessibility to the amine of G, which may account for its enrichment as a dual donor.

**Figure 3.**
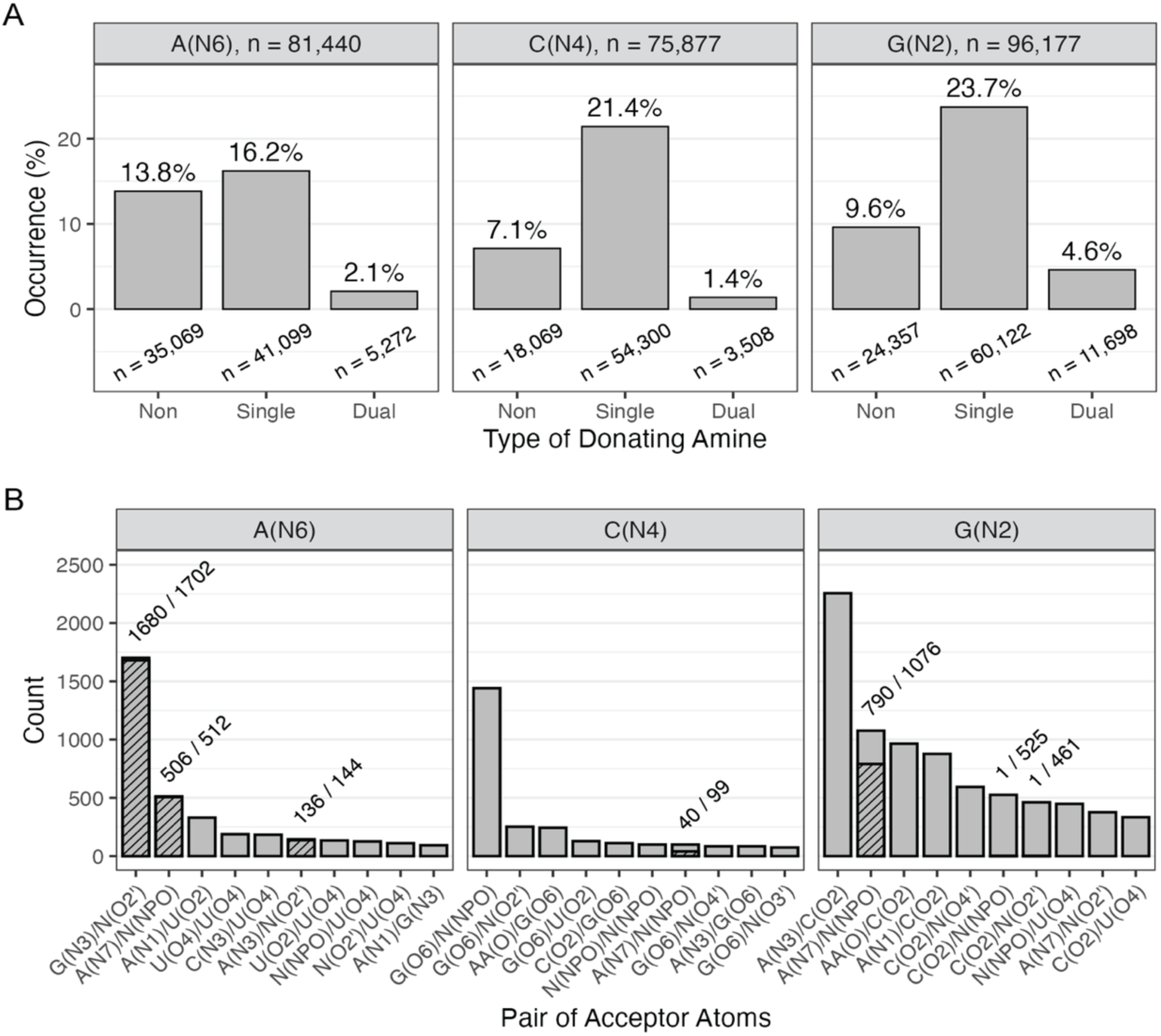
Prevalence of H-bonding interactions amongst amines. (*A*) Percentage of non-, single-, and dual-donating amines amongst all A’s, C’s, and G’s. All columns sum to 100%. The number of amines surveyed are provided above the plots and the number belonging to each donor type are provided below its respective column. (*B*) Number of acceptor pairs that are engaged with dual-donating amines. Hatched portions of columns represent the number of acceptor pairs that are part of the same residue, and the numbers above these columns indicate the fraction of pairs belonging to the same residue. When the acceptor is a backbone atom, any canonical RNA residue is denoted with “N” and amino acid residue with “AA”. The top 10 acceptor pairs per nucleobase are provided. Dual-donating amines with more than two H-bonds were handled as described in the Methods. Select examples of some of these interactions are depicted in Figure 4 and Supplemental Figure S1.

Moving to single-donating amines, these are the largest proportion of the distribution at 61.3% (16.2% A + 21.4% C + 23.7% G), likely a consequence of the abundance of canonical base pairing interactions. The enrichment of C and G over A might reflect the bias of the structural biology community to study strongly folding RNAs such as rRNA. Finally, non-donating amines account for the remaining 30.5% (13.8% A + 7.1% C + 9.6% G) of all amines, with an enrichment in A, which may reflect the above biases.

We turn our focus to the identity of the atom pairs that accept the two H-bonds from the dual-donating amines. Starting with dual-donating A(N6), it is striking that the three top acceptor pairs do not involve U(O4), the acceptor in a canonical AU base pair (Figure 3B), which indicates that most A dual-donating amines do not simply recruit a second H-bonding acceptor to a canonical AU base pair. The dominant acceptor pair is G(N3)/N(O2′), and in 1,680 out of 1,702 cases these two acceptors belong to the same G, an example of which is the sheared GA base pair provided in Figure 4A. Sheared GA base pairs form when the Hoogsteen edge of the A interacts with the sugar edge of the G, with the dual-donating amine providing two H-bonds to this interaction (Jang et al. 2004). The sheared GA dominates the G(N3)/N(O2′) acceptor pair category, comprising 1,518 of 1,702 instances, consistent with the high abundance of this non-canonical base pair (Olson et al. 2019). All instances of the interactions depicted in Figure 4, from all relevant PDB entries, are provided within CSV and Python files that are available from the Supplemental Material (listed in Supplemental Table S1) and can be reviewed visually by running the Python files within a PyMOL session (see Methods).

**Figure 4.**
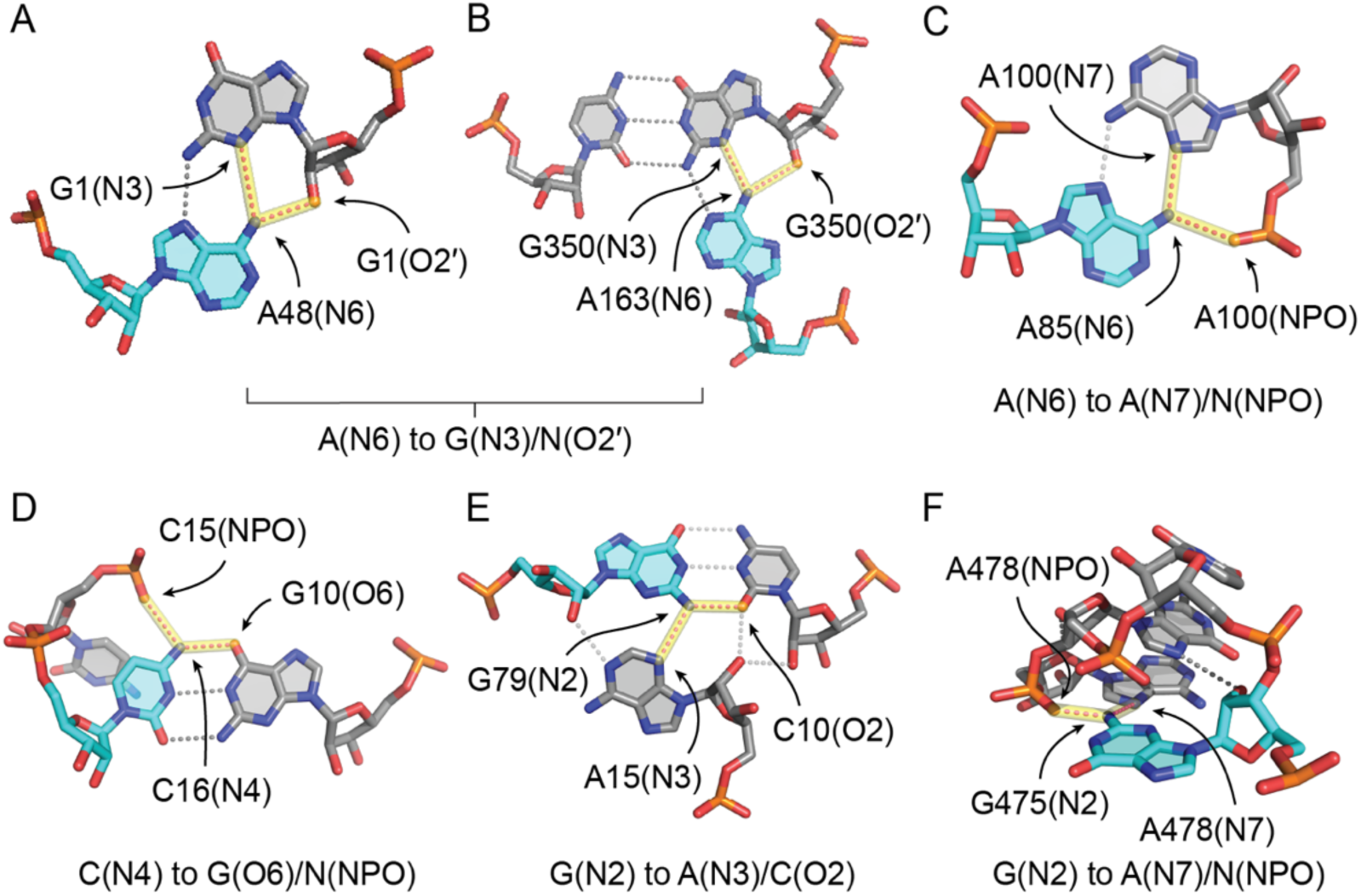
Common acceptors for dual-donating amines. (*A-B*) Adenine dual-donating amines interacting with the N3 and O2′ of the same G forming (*A*) a sheared GA base pair involving A48 from chain A of PDB ID 3FS0 (Pitt and Ferré-D’Amaré 2009) and (*B*) a WC/H A-minor motif involving A163 from chain L50 of PDB ID 7QCA (McLaren et al. 2023). (*C*) A85(N6) dual donates to the N7 and NPO of the same A, where A85 is from chain 2 of PDB ID 9AXU (Gemin et al. 2024). (*D*) C16(N4) dual donates to a G(O6) in a canonical GC base pair and an NPO from a different residue, where C16 is from chain 6 of PDB ID 7B9V (Wilkinson et al. 2021). (*E*) G79(N2) dual donates to a C(O2) in a canonical GC base pair and an A(N3), forming a traditional A-minor motif. G79 is from chain A5 of PDB ID 8A22 (Tobiasson et al. 2022). (*F*) G475(N2) dual donates to the N7 and NPO of the same A within a GNRA tetraloop, where G475 is from chain AA of PDB ID 6ERI (Perez Boerema et al. 2018). In all six panels, magenta dots highlighted in yellow depict H-bonds from dual-donating amines, and grey dots represent other H-bonds. The subtitle under each panel corresponds to a bar depicted in Figure 3B. Instances of these dual-donor-to-acceptor-pair interactions are provided within CSV and Python files available in the Supplemental Material, which are listed in Supplemental Table S1.

Another example of the G(N3)/N(O2′) accepting pair from A is in the A-minor motif (Figure 4B). This is one of the most prevalent motifs in structured RNAs and involves the interaction of an A with the minor groove of another pair (Baulin 2021). Intriguingly, J. Steitz and colleagues recently described a new subclass of A-minor motif that they coined the “WC/H A-minor” motif (Torabi et al. 2021), which this example fits into. This newly defined subclass involves the WCF and/or Hoogsteen edge of A interacting with the minor groove of a base pair. Thus, any of the above G’s that also form a canonical GC base pair, would result in a base triple that would be part of a WC/H A-minor motif. We identified 40 instances of this motif, expanding the examples articulated by the Steitz group. In contrast to the sheared GA base pairs where G(N2) donates to A(N7), most of these WC/H A-minor instances (31 out of the 40) involves G(N2) donating to A(N1) since the A(N7) is swung away from the G(N2). Of note, the G(N2) is itself a dual-donating amine in nearly all cases (38 out of 40).

After the G(N3)/N(O2′) accepter pair, the next most common acceptor pair for A(N6) is A(N7)/N(NPO), where NPO stands for non-bridging phosphoryl oxygen, with a count of 512 (Figure 3B). In nearly all of these occurrences (506 out of 512), the N7 and NPO acceptors belong to the same residue, making them interactions between two A’s. Furthermore, the neighboring A donates an A(N6)-to-A(N7) H-bond to the A bearing the dual-donating amine in most cases (458 out of 506) (Figure 4C), resulting in a trans base pair involving the Hoogsteen edge of both A’s and at least one NPO. The large negative charge of the NPO acceptor likely plays a key role in stabilizing this interaction (see Discussion). The remainder of the acceptor pairs for A(N6) occurred considerably less frequently, with counts of 330 or less.

Moving to C, the top five acceptor pairs contain G(O6) (Figure 3B), the acceptor in canonical base pairs, deviating from what was found for A(N6). The most common acceptor pair for C(N4) is G(O6)/N(NPO), with a count of 1,441, and these two acceptors never belong to the same residue (Figure 3B). Given the negative charge on the phosphate, one would expect a significant electrostatic component for this attractive interaction. As depicted in Figure 4D, the parent C of the dual-donating amine forms a canonical base pair with the G that accepts the H-bond from C(N4), which occurs in most cases (1,400 out of 1,441). The remainder of the acceptor pairs for C(N4) drop quickly in count.

Considering G, the interactions of its amine differ from those of A and C in that the various acceptor pairs do not drop in count nearly as quickly (Figure 3B), indicating that G’s amine engages in a more diverse set of dual-donating interactions. The dominant acceptor pair for G(N2), in 2,256 instances, consists of A(N3)/C(O2), where the latter is the WCF H-bonding partner of G(N2) (Figure 3B). Within the majority of the cases (2,178 out of 2,256), the G forms a canonical pair with the C and the sugar edge of A interacts with the minor groove of the GC pair, making this a traditional A-minor motif (Figure 4E). The second most common acceptor pair for G(N2) is A(N7)/N(NPO) (Figure 3B).

Of the 1,076 amines in this category, 790 donate to acceptors that belong to the same A residue. Visual inspection of a sample of this subset reveals tetraloops and pentaloops contributing to this dual-donating interaction, where the G and A make up the first and last members of the loop, respectively (Figure 4F).

Finally, we note that for both C(N4) and G(N2), the third most prevalent acceptor pair consists of an amino acid backbone carbonyl oxygen, AA(O), and the acceptor relevant to canonical base pairing (i.e., G(O6) and C(O2), respectively), as depicted in Supplemental Figure S1. For dual donation by C(N4), the AA(O) is in the major groove, while for G(N2), it is in the minor groove, making the dual-donating amine versatile in positioning. Like the phosphate backbone described above, the backbone carbonyl oxygen has significant negative charge, which could strengthen these H-bonds (see Discussion). Notably, no amino acid acceptor was present in the top ten acceptor pairs for A(N6). Evidently, dual-donating amines belonging to C and G play a greater role in protein binding in this context. All instances of the interactions depicted in Supplemental Figure S1 are provided within Python files (for visualization using PyMOL) and CSV files that are available from the Supplemental Material (listed in Supplemental Table S1).

*Dual-Donating Amines Tend to Be Buried in Solvent Inaccessible and Dense Regions* Owing to long-range and often non-canonical interactions, highly structured RNAs are often globular-like, in which some residues reside closer to the interior. We were curious whether dual-donating amines could drive such compaction. To test this possibility, we measured the solvent accessibility of the exocyclic amines and the density of the surrounding atoms.

We began assessing burial by considering solvent accessible surface area (SASA), which can measure exposure of a residue or atom to solvent. To measure SASA, we used the Biopython suite of tools, which implements an algorithm based on that developed by Shrake and Rupley (1973). In all cases, we measured the SASA of the nitrogen of the amine, where no hydrogens were present in the structure. We found that the median SASA values for the amine nitrogens of A, C, and G dropped monotonically along the series non-, single-, and dual-donating, where it fell to 0 Å^2^, indicating that dual donors are fully buried (Figure 5A and Supplemental Figure S2A). Both A’s and G’s exhibited similar SASA values for their amine nitrogens within the non- and single- donating categories, with respective values of 9.8 Å^2^ and 4.4 Å^2^ for A(N6) and 9.8 Å^2^ and 5.5 Å^2^ for G(N2). In contrast, the median SASAs for the non- and single-donating categories were elevated for C(N4), with values of 14.2 Å^2^ and 8.7 Å^2^, respectively, which may be due to the absence of the imidazole ring which can stack and whose N7 can assist in burial. Despite higher values for the non- and single-donating categories, the median SASA for dual-donating C(N4) is still 0 Å^2^, reinforcing the point that dual-donating amines are in the interior of an RNA. Notably, the median SASA values of 0 Å^2^ for dual- donating amine nitrogens from all three bases indicates that there is crowding from the atoms of other residues not just along the edges of the donating base, but above and below it as well. The importance of atoms above and below a dual-donating amine was confirmed by omitting all atoms other than the residues involved in a dual-donating interaction—we chose the central C(N4) within the base triple depicted in Figure 5B—and finding a sizeable SASA (10.9 Å^2^ in this example). Finally, we do note that a slight fraction of dual donors has non-zero, albeit very small, SASA values (Supplemental Figure S2B).

**Figure 5.**
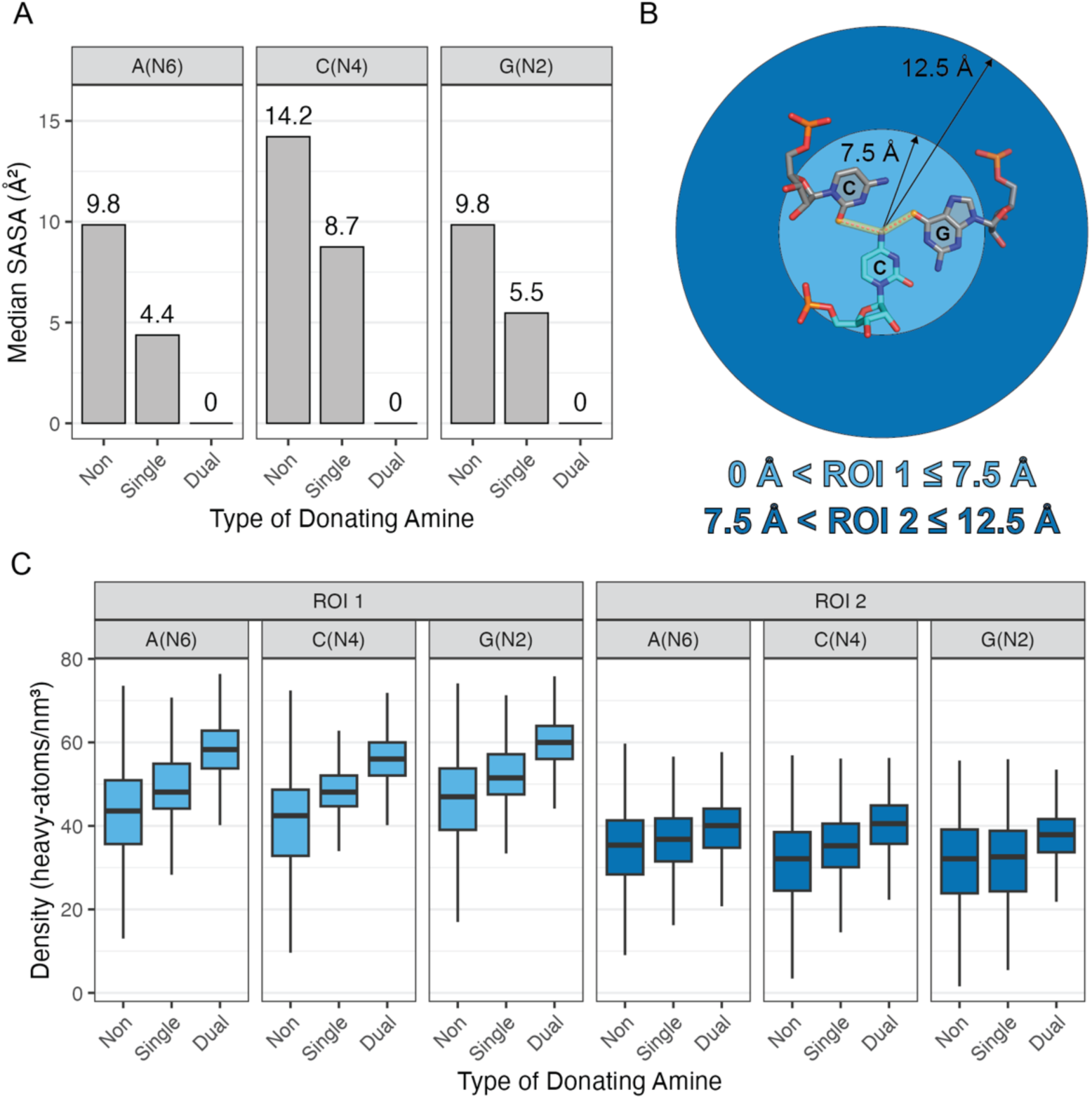
Amine solvent accessibility and surrounding density. (*A*) Median SASA values for the nitrogen atoms of non-, single-, and dual-donating amines for each type of nucleobase. Corresponding box plots are provided in Supplemental Figure S2A. (*B*) Diagram depicting the defined first region of interest (ROI 1 in light blue) and the second region of interest (ROI 2 in dark blue). The regions are centered around the amine of interest, which in this example is C1092(N4) from chain C of PDB ID 1MMS (Wimberly et al. 1999). (*C*) Density of heavy atoms within ROI 1 and ROI 2. All adjusted p-values for heavy atom density are less than 1×10^-5^ for the differences observed in the distributions within each of the six subplots (see Methods). Outliers were omitted from these plots but are included in Supplemental Figure S2C.

We next assessed the extent of burial of the dual-donating amines by measuring the density of surrounding heavy (i.e., non-H) atoms. This quantity was measured in two regions of interest (ROIs), an inner one within 7.5 Å of the amine, termed “ROI 1”, and an outer one between 7.5 Å and 12.5 Å of the amine, termed “ROI 2” (Figure 5B). In addition to heavy atoms from RNA, we included atoms from proteins and other polymers. However, heavy atoms from water, inorganic species, and organic molecules (i.e., ligands) were not included. The 7.5 Å cut-on for ROI 2 was chosen as it removes atoms in direct contact with the amine. It also defines the boundary of the nucleobases in a triple base pair with the dual-donating amine at the center, as depicted in Figure 5B.

For the ROI 1 densities, the medians for non-, single-, and dual-donating A amines increased monotonically, with values of 44, 48, and 58 atoms/nm^3^, respectively, as depicted in the light blue box plots in Figure 5C and Supplemental Figure S2C (outliers included). The medians for C and G showed the same trend, with values of 42, 48, and 56 atoms/nm^3^ and 47, 51, and 60 atoms/nm^3^, respectively. Moreover, the magnitude of the increase in density was greater going from single- to dual-donating (changes of 10.2, 7.9, and 8.5 atoms/nm^3^ for A’s, C’s, and G’s, respectively) when compared with going from non- to single-donating (changes of 4.5, 5.7, and 4.5 atoms/nm^3^). Additionally, the distributions for the dual-donating amines were tighter, with average IQR across all three nucleobases decreased by 10% relative to single-donors and by 46% relative to non-donors. The narrowness of the distributions suggests that higher density is a consistent property of dual-donating amines.

For ROI 2, the median densities also increased monotonically upon going from non- to single- to dual-donating amines for all three nucleobases, albeit with a more moderate incline, as depicted in the dark blue box plots in Figure 5C and Supplemental Figure S2C (outliers included). Moreover, the magnitude of the increase in going from single- to dual-donating remained greater than that for non- to single-donating, and the IQRs again tightened as the amine progresses from non- to single- to dual-donating. This tightening of IQR is especially prominent for G(N2), where the dual category IQR (8.0 atoms/nm^3^) was nearly half that of the single category IQR (14.5 atoms/nm^3^). It is remarkable that higher atom density and tighter IQR persist in amine-distal regions over 12 Å away.

### Glycoside Bond Dihedrals Vary Depending on Amine H-Bond Donation

Owing to rotations available at the glycosidic bond and six sugar-phosphate backbone dihedral angles, RNA is an extraordinarily flexible polymer. However, because it engages in multiple interactions, dual-donating amines have the potential to reduce the conformational entropy loss upon RNA folding, which might enhance certain RNA conformations. We therefore investigated whether donating amines might show preference for rare conformations in their glycosidic bonds.

The *χ* dihedral reflects a nucleotide’s conformation about its glycosidic bond and is typically categorized as being *anti* or *syn*, in other words, with the base away from or over the sugar, respectively. Our lab previously conducted a structural survey of functional RNAs for nucleobases in the rare *syn* conformation (Sokoloski et al. 2011). An amine engaged in dual donation could, in principle, help stabilize this energetically unfavorable conformation. To test for this, we measured the *χ* dihedrals of residues bearing non-, single-, and dual-donating amines.

A histogram for all bases revealed a major and a minor conformation, with the latter mostly corresponding to the *syn* orientation (Figure 6A), where *syn* nucleobases are defined as having *χ* dihedrals ranging from -90° to 90° (Bloomfield et al. 2000). We then prepared nine density plots, one for each nucleobase and amine donation type. This was done across the entire -180° to 180° range (Supplemental Figure S3), with just the -20° to 130° range, corresponding to the minor conformation, displayed in Figure 6B.

**Figure 6.**
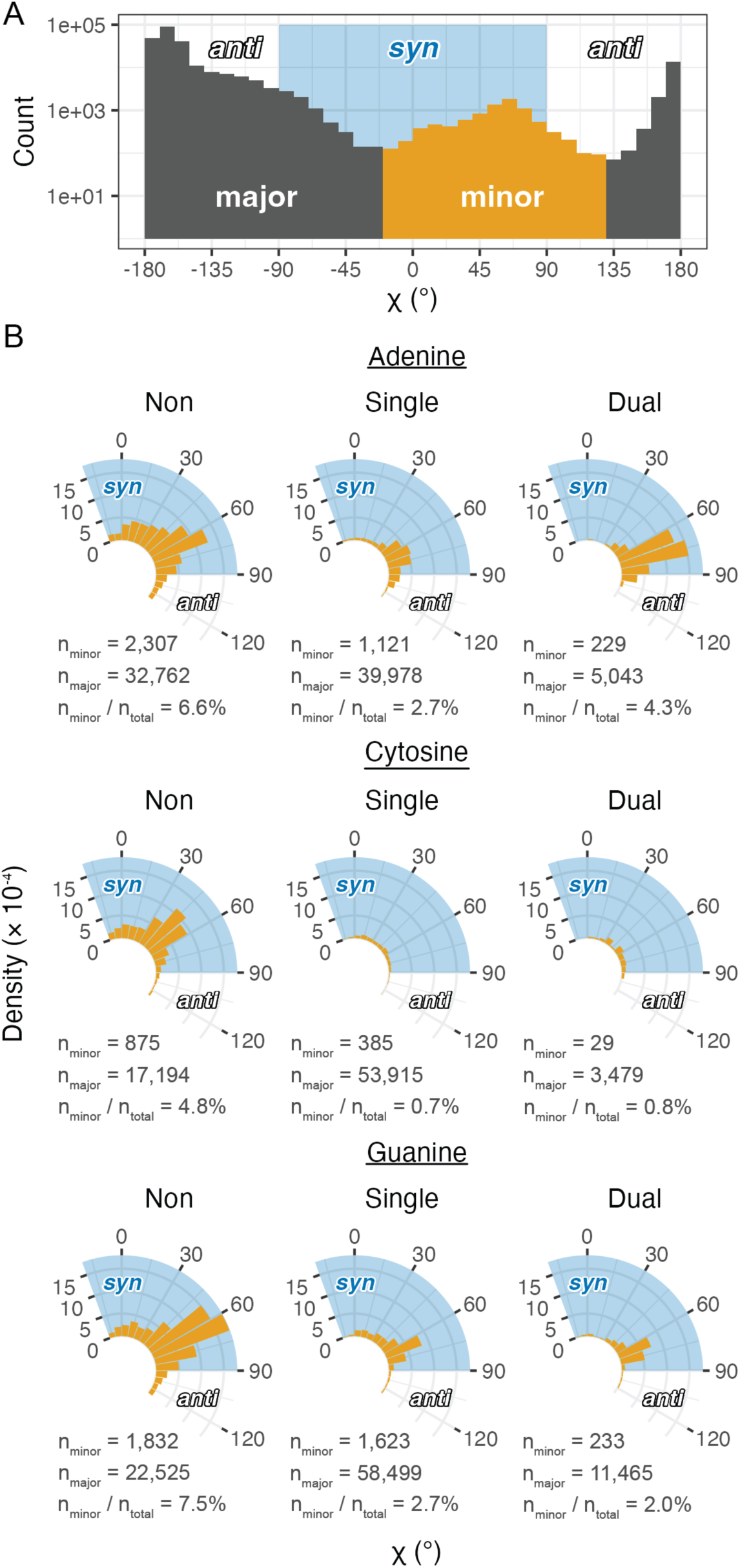
Analysis of *χ* dihedrals of the minor population. (*A*) Histogram (logarithmic scale) spanning the entire *χ* dihedral range. A minor population (orange bars) is found and is primarily in the *syn* conformation (blue shading). (*B*) Radial density plots of the minor population *χ* dihedrals separated into non-, single-, and dual-donating amines. Densities are based on calculations of the full dihedral range presented in Supplemental Figure S3. The number of residues in the minor and major conformations are provided under each plot for all nucleobase and donating amine types, along with the corresponding percentages of residues that occupy the minor conformation. The blue shaded regions correspond to nucleobases that adopt the *syn* conformation, which extends beyond the lower bound of these plots to -90°.

We first focus on the minor conformation given the functional importance of *syn* nucleobases (Sokoloski et al. 2011). A’s with dual-donating amines are enriched when compared to those with single-donors (4.3% versus 2.7%) (Figure 6B). For C, however, both single- and dual-donating amines are depleted (0.7% and 0.8%). It is well known that pyrimidines adopt the *syn* conformation less frequently than purines owing to clash of the O2 with the ribose sugar (Saenger 1973; Sundaralingam 1975; Saenger 1984; Sokoloski et al. 2011). Lastly, for G, there appears to be a slight preference for single- donating amines over dual donors (2.7% versus 2.0%), and the dual-donating amines are intermediate in population to A and C. Relative to A’s, the lower population of G’s with dual-donating amines in the minor conformation may be due a paucity of acceptors on the top face of the sugar and/or to steric clash of the G(N2). Across all three nucleobases, any type of H-bond donation by the amine disfavors the minor conformation, indicated by the relatively higher populations for the non-donors (6.6% for A, 4.8% for C, and 7.5% for G). This is likely because the major conformation exposes the amine to a wider array of potential acceptors and because WCF base pairs are common and adopt the *anti* conformation.

Focusing on the major conformation (Supplemental Figure S3), we note that amine H-bond donation narrows the distribution of the *anti* conformation for all three bases, albeit dual-donation pushes A back towards a somewhat broader distribution. Apparently, single- and dual-donation restricts the conformational heterogeneity of the nucleosides.

### Pseudo-Torsional Analysis Reveals Dual-Donating Amine Enrichment in S-motifs

To further assess the enrichment of dual-donating amines in select conformations, we next considered conformations along the backbone. For ease of analysis, the six covalent bonds linking the backbone together were reduced to two virtual bonds, one involving P_!_ − C4’_!_ and another involving C4’_!_ − P_!"#_ , as described previously (Olson 1976; Duarte et al. 2003). These virtual bonds give rise to two pseudo-torsion angles, η and θ, C4’_!$#_ − P_!_ − C4’_!_ − P_!"#_ and P_!_ − C4’_!_ − P_!"#_ − C4’_!"#_ , respectively, where *i* represents the residue of interest. This reduction allows Ramachandran-like plots to be constructed, in which different RNA motifs tend to occupy different regions; for instance, A-form helices exhibit η and θ values near 170° and 220°, respectively (Keating et al. 2011).

In an effort to identify any RNA motifs unique to dual-donating amines, we plotted heatmaps of the pseudo-torsion angles for non-, single-, and dual-donating amines (Figure 7A-C). Pseudo-torsion conformations corresponding to A-form helices, with values of 145° > η > 190° and 190° > θ > 245°, dominate the plots as previously reported (Duarte and Pyle 1998; Humphris-Narayanan and Pyle 2012; Grille et al. 2023). For dual- donating amines, two locations with relatively high counts also emerged: one near η of 55° and θ of 165° (Location 1) and another near η of 310° and θ of 30° (Location 2) (Figure 7C). For non- and single-donating amines, the same two regions are not highly enriched (Figure 7A and 7B).

**Figure 7.**
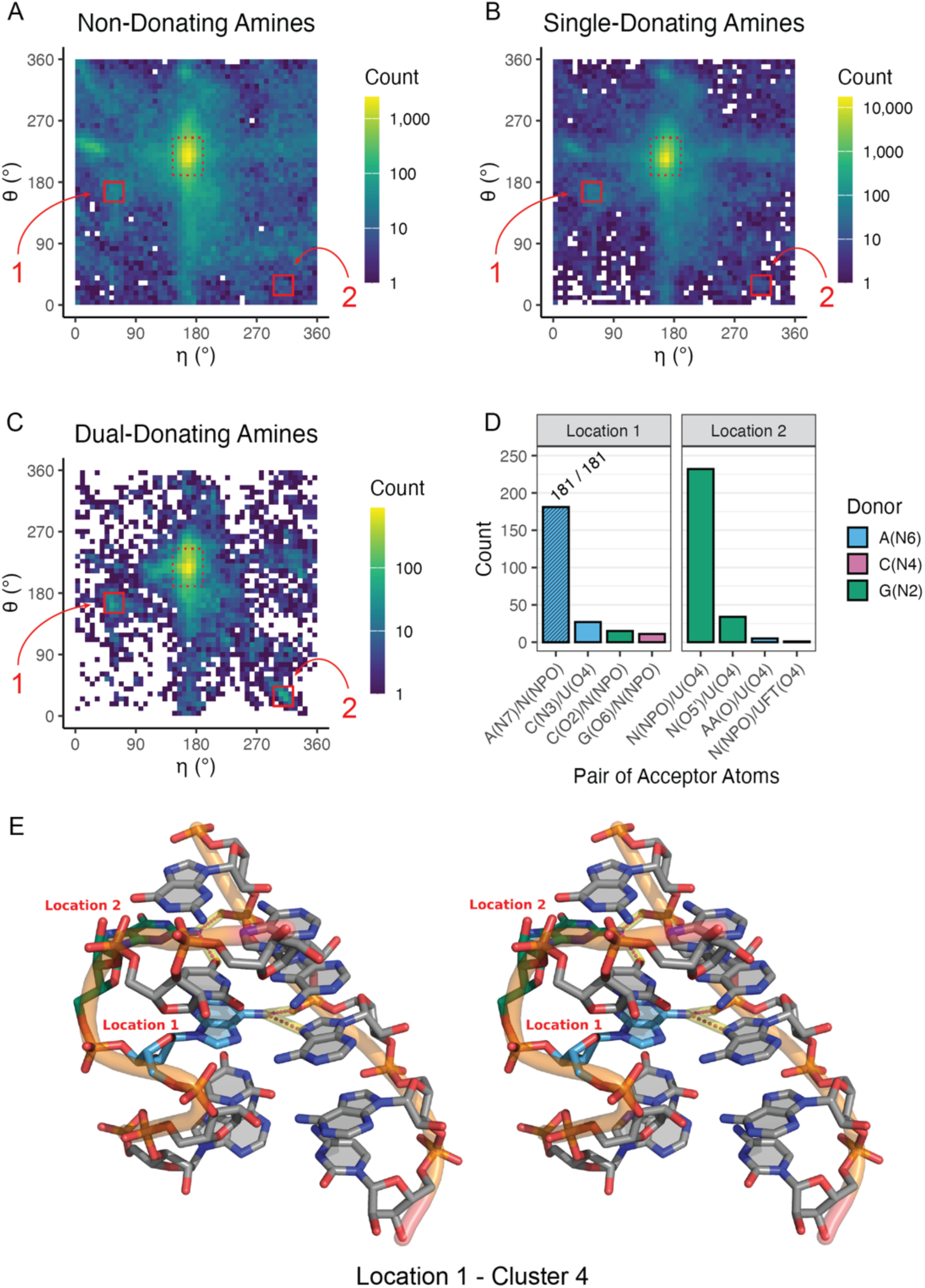
Analysis of pseudo-torsion angles reveals key dual-donating amines involved in S-motifs. (*A-C*) Heatmaps depicting pseudo-torsion angles of residues bearing (*A*) non-, (*B*) single-, and (*C*) dual-donating amines. The color scale is logarithmic to accommodate the wide-range of data. Locations of interest, “Location 1” and “Location 2”, are boxed in red. The residues within Location 1 and Location 2 of (*C*) are provided within CSV and Python files available in the Supplemental Material, which are listed in Supplemental Table S1. (*D*) Number of acceptor pairs that are engaged with dual-donating amines per nucleobase for Location 1 and Location 2. Symbols “N” and “AA” are used to denote any canonical RNA or amino acid residue, respectively, when the acceptor is a backbone atom. Only the top four acceptor pairs are shown for each Location. (*E*) Cross-eye stereoview of the representative structure (PDB ID 6ZU5) from the dominant cluster (C4) of the Location 1 cluster analysis (Ehrenbolger et al. 2020). An S-motif is depicted with an A dual-donating amine from Location 1 (blue) and a G dual-donating amine from Location 2 (green). Magenta dots highlighted in yellow depict H-bonds from the two dual-donating amines. A similar depiction corresponding to the Location 2 cluster analysis can be found in Supplemental Figure S6A.

To determine the dual donors and their acceptors in Location 1 and Location 2, bar charts were constructed (Figure 7D). For Location 1, which had η values ranging from 43 to 72° and θ values ranging from 151 to 180°, there was a single dominant interaction, that of A(N6) donating to A(N7)/N(NPO), with a count of 181. This matched the second most prevalent acceptor pair for all A dual-donating amines (Figures 3B) where the majority of such pairs involved a trans base pair along the Hoogsteen edges of two A’s (Figure 4C). For Location 2, which had η values ranging from 295 to 324° and θ values ranging from 14 to 43°, there was again a single dominant interaction, here of G(N2) donating to N(NPO)/U(O4), with a count of 232. This corresponds to the eighth most prevalent acceptor pair for all G dual-donating amines (Figure 3B). It is notable that the second most prevalent interaction, with a count of 34, was like the first, with the only difference being that the first acceptor is a bridging, rather than a non-bridging, phosphoryl oxygen. Further analysis revealed that 83 residues in Location 1 were connected to the 5′-end of a residue from Location 2; thus, the residues from Location 1 and Location 2 are sometimes adjacent in sequence. These 83 instances, along with all other residues bearing dual-donating amines and corresponding to Location 1 or Location 2, are provided within Python files (for visualization using PyMOL) and CSV files that are available from the Supplemental Material (files listed in Supplemental Table S1).

After considering plots similar to Figure 7A-D, we manually inspected a variety of structures contributing to Location 1 and Location 2 to better understand the context of these dual-donating amines. It became clear that many of them belong to S-motifs, which make up part of the sarcin/ricin loop in ribosomal RNA, and we found that the corresponding η and θ values are close to previously reported values for these motifs (Correll et al. 2003; Duarte et al. 2003). To identify a representative structure for each Location, we carried out a clustering analysis (see Methods). We first extended the sequence in both directions along the strand containing the residue of interest and then repeated this for its base pairing partner, resulting in a 6x5 nucleotide fragment. This process was conducted on all residues for the major acceptor pair for Location 1 (n = 181) and all residues for the two major acceptor pairs for Location 2 (n = 266). The fragments in Location 1 were superposed with each other, as were the fragments in Location 2. Cut- offs of 3.40 Å for Location 1 and 4.10 Å for Location 2 were chosen on the basis of the resulting Silhouette scores (Supplemental Figure S4). The resulting dendrogram had four clusters for Location 1, with the dominant cluster (C4) containing 97 of the 181 fragments, as well as four clusters for Location 2, with the dominant cluster (C3) containing 158 of the 266 fragments (Supplemental Figure S5). The representative structure for each cluster was the structure with the smallest RMSD to the cluster’s average structure (see Methods).

As expected, the representative structure from the dominant cluster for Location 1 (Figures 7E) and the representative structure from the dominant cluster for Location 2 (Supporting Figure S6A) exhibit S-motifs. These examples display the characteristic “S” shape along the backbone, depicted as the left-most strand in these two figures. Notably, in each example, the dual-donating amine of G in Location 2 (depicted in green) is bulged, while the dual-donating amine of A in Location 1 (depicted in blue) forms a trans base pair with another A. It is striking to note the network of other hydrogen bonds and stacks made possible by these dual-donating bases. Notably, the sequence 5′-AGU-3′ in the S-motif has the U stacking on the A, where the G is in Location 2 and the A in Location 1, and one of the acceptors of the dual-donating amine of G is the O4 of the U. Out of the eight clusters identified, six exhibited S-motifs: C1, C2, and C4 for Location 1 and C1, C3, and C4 for Location 2 (Figure 7E and Supplemental Figure S6), with the two missing clusters having relatively few fragments. Altogether, an analysis of pseudo-torsion angles followed by clustering of fragments revealed that dual-donating amines are abundant in S-motifs at two distinct positions, with both dual donors sometimes occurring in the same structure.

### Key RNA Motifs in Functional RNAs Contain Numerous Dual-Donating Amines

Dual-donating amines play essential roles in functional RNAs. Indeed, we found multiple dual-donating amines in a number of key RNA motifs. To appreciate their importance to RNA structure, we feature three such motifs here (Figure 8, Supplemental Figure S7 is stereo). The 11-nucleotide GNRA tetraloop receptor contains an internal loop that binds a GNRA tetraloop via tertiary interactions (Juneau et al. 2001; Butcher and Pyle 2011). An example from the self-splicing group I intron reveals five dual-donating amines within the confines of this relatively small structure (Figure 8A, Supplemental Figure S7A is stereo). Two of the dual-donating amines correspond to the two most prevalent acceptor pair categories for G(N2) (Figure 3B): one G(N2) is in the receptor and forms an A-minor motif with the GAAA tetraloop, donating to A(N3)/C(O2) (Figure 4E), while the other G(N2) is the first residue in the GAAA tetraloop, donating to A(N7)/N(NPO) (Figure 4F); notably, a single A interacts with both of these G’s. Additionally, one of the A dual-donating amines from the receptor H-bonds with an A(N1) from the tetraloop and a U(O2) from the receptor, corresponding to the third most prevalent acceptor pair for A(N6) in Figure 3B. Remarkably, most residues in the receptor and the bound GAAA tetraloop are involved in dual-donating amine interactions, either as a donor, an acceptor, or both, emphasizing the prevalence of these interactions in tertiary structure.

**Figure 8.**
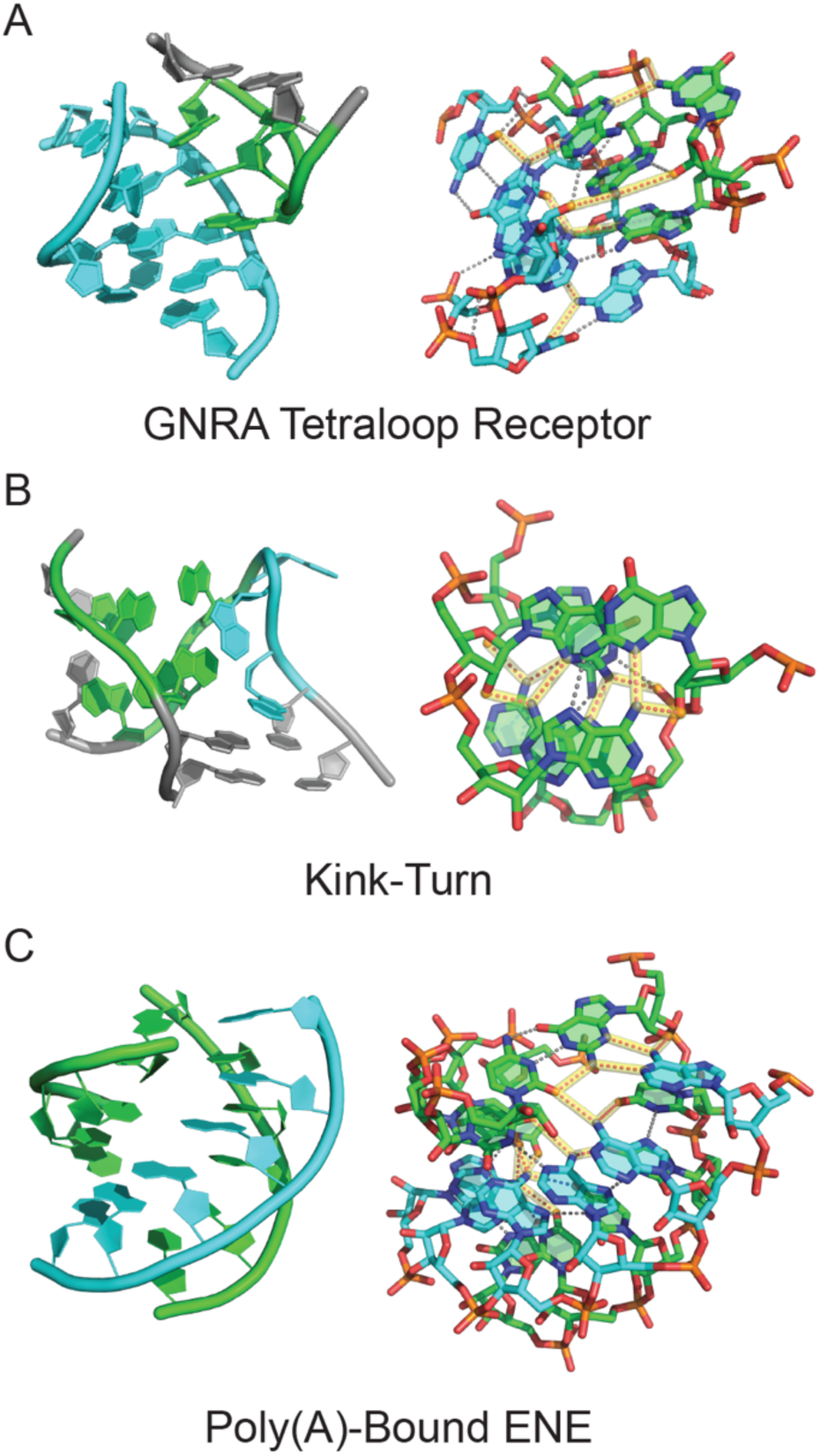
Structured RNAs enriched in dual-donating amines. (*A*) GNRA tetraloop receptor (cyan) bound to a GAAA tetraloop (green) from the P4-P6 domain of the group I intron with five dual- donating amines (PDB ID 1HR2). (*B*) Kink-turn from the SAM riboswitch with a 3-nucleotide bulge (cyan) and series of sheared GA base pairs (green) with four dual-donating amines (PDB ID 2GIS). (*C*) Poly(A) strand (cyan) bound to the U-rich internal loop and GC closing base pair of an ENE (green) from a rice transposase mRNA with seven dual-donating amines (PDB ID 7JNH). For all panels, left is a cartoon representation, and right is a stick representation. Magenta dots highlighted in yellow depict H-bonds from dual-donating amines, and grey dots represent other H-bonds. Cross-eye stereoviews are provided in Supplemental Figure S7.

Next, we feature the kink-turn motif in the S-adenosylmethionine (SAM) riboswitch, which involves a 3-nucleotide bulge loop flanked on one side by three tandem sheared GA base pairs (Montange and Batey 2006; Butcher and Pyle 2011; Lilley 2012) (Figure 8B, Supplemental Figure S7B is stereo). As the name suggests, the kink-turn motif results in a severe bend between the axes of the helices flanking the bulged loop. A crystal structure of the SAM riboswitch depicts four dual-donating amines in the three tandem sheared GA base pairs, with the central GA pair having two dual-donating amines and the terminal GA pairs each having one dual-donating amine. The A amines of these sheared GA pairs dual donate to G(N3)/N(O2′), which corresponds to the most prevalent acceptor pair category for A(N6) (Figures 3B and 4A). These non-canonical base pairs hold the phosphate backbone of the opposing strands closer together, with interstrand phosphate distances of 12.9 to 16.0 Å versus 18 to 21 Å for a normal RNA helix (Supplemental Figure S8) (Pande and Nilsson 2008). With four dual-donating amines, the density of these amines in the kink-turn is similar to that in the tetraloop receptor.

Finally, we present a structure of a poly(A)-bound element for nuclear expression (ENE) (Torabi et al. 2021). The poly(A) tail of an ENE-containing transcript binds the U-rich internal loop of the ENE to form an RNA triplex structure, which, for example, increases the half-life of an oncogenic Kaposi’s sarcoma-associated herpesvirus (KSHV) transcript (Conrad and Steitz 2005; Mitton-Fry et al. 2010). Two ENEs—an upper and a lower—from a rice transposase mRNA are depicted in a crystal structure with a bound poly(A), revealing many interactions involving dual-donating amines (Figure 8C, Supplemental Figure S7C is stereo). Here, we focus on the U-rich internal loop and its closing GC base pair of the upper ENE, complexed with the poly(A). Seven dual-donating amines span across this segment. Four contribute to UAU triplets of the U-rich internal loop, where the A(N6) dual-donates to a U(O4)/U(O4) acceptor pair, the fourth most prevalent acceptor pair for A(N6) (Figure 3B). The other three dual-donating amines participate in a quintuple-base interaction that is at the top of the U-rich internal loop and involves the closing GC base pair. This quintuple-base interaction contains a WC/H A-minor motif which corresponds to the top acceptor pair for A(N6) (Figures 3B and 4B). In sum, the GNRA tetraloop receptor, kink-turn motif, and poly(A)-bound ENE are central to catalytic introns, riboswitches, and transcripts in cancer-causing viruses and display a remarkably high density of dual-donating amines. Clearly, dual-donating amines can play a critical role in stabilizing and compacting the tertiary structure of functional RNAs.

## Discussion

We developed a cheminformatic pipeline for assessing the prevalence and context of dual-donating amines in RNA structure. High resolution structures of RNAs, curated by the Representative Sets of RNA 3D Structures, were inspected for hydrogen bonds from each of the two hydrogens of exocyclic amines, using a set of distance and angle criteria determined herein. We then classified nucleobase amines as non-, single-, or dual- donating and performed further analysis, including the consideration of non-canonical geometries. Importantly, this approach is reliant on experimental RNA structures deposited in the PDB; as such, our findings do not reflect any special characteristics of under- or non-represented RNAs.

Our study uncovered an enrichment of dual-donating amines in diverse base pairs including 1) trans AA pairs, which contribute to S-motifs, 2) A-minor interactions, which occur in GNRA tetraloop receptors, 3) sheared GA pairs, which are present in kink-turns, and 4) WC/H A-minor interactions, which occur in some ENE motifs (Figures 4, 7, and 8). We also identified dual-donating amines serving key roles in engaging with amino acids in both the major and minor grooves (Supplemental Figure S1). Additionally, we discovered that the dual-donating amine motif is largely buried, with persistent high atom density and very limited solvent accessibility (Figure 5 and Supplemental Figure S2). Remarkably, the median SASAs for all dual-donor categories were 0 Å^2^ despite the calculations being done without hydrogens.

One reason why dual-donating amines may play an outsized role in RNA structure is that RNA has both an anionic backbone and is replete in acceptors (Figure 1). The 5- fold excess of H-bond acceptors over donors makes RNA largely self-avoidant with its phosphate backbone. Indeed, when multiple H-bond donors are present in a single molecule, such as urea with two amines, RNA will unfold (Shelton et al. 1999; Lambert and Draper 2012; Jaganade et al. 2019; Raghunathan et al. 2020). The abundance of H-bond acceptors makes a dual-donating amine capable of stitching together local RNA structures, as evidenced by turn motifs, including the kink-turn and S-motif, and forming long-range interactions, such as between the GNRA tetraloop and its receptor, which compact the RNA.

The strength of base pairing may be enhanced by the dual-donating amine, which it can accomplish by reducing the conformational entropy loss for folding and strengthening H-bonding. Forming H-bonds incurs an entropic penalty of loss of motion. Since the amine loses that entropy once in forming the first H-bond, it may not have to lose it again in forming the second H-bond, providing a reduction in conformational entropy loss, favoring folding. There are many diverse dual-donating amine motifs that could aid folding (Figure 4). Some of these motifs may also have strengthened H-bonding, particularly those found in Figures 4C, D, and F and Supplemental Figure S1. In these cases, the dual-donating amine donates to either a phosphate or a carbonyl of the peptide backbone. In two of these cases (Figure 4D and Supplemental Figure S1A), the C of a canonical GC base pair donates to either a phosphate or the carbonyl of the peptide backbone. Notably, the phosphate and carbonyl groups have estimated atomic charges of –0.78 and –0.57, respectively (Cornell et al. 1996). Interactions with these anionic species should favor stronger electron donation by the amine of the C into the ring system. This would result in the development of greater positive charge on the N4 and negative charge on the N3 and O2. The charge development on these three atoms should move their p*K*a’s towards neutrality, which should in turn strengthen the H-bond to the G due to better p*K*a matching (Herschlag and Pinney 2018). Strong electron donation into the ring system may also perturb other chemical properties of the nucleobase, which could have implications for covalent modifications and chemical probing. Finally, the newfound knowledge of how dual-donating amines contribute to important structural motifs could prove useful in RNA 3D structure prediction.

## Methods

### Experimental Structures and Representative Dataset

*(i) Reducing redundancy in RNA-containing structures.* While the PDB includes many RNA-containing entries, a number of these structures are highly similar to one another. For instance, a search on rcsb.org for structures containing a Polymer Entity Description matching “28S ribosomal RNA” or “28S rRNA” and the Scientific Name of the Source Organism matching “*Homo sapiens*” returned 179 results (as of May 28, 2025). The Representative Sets of RNA 3D Structures offers a well-documented and traceable solution to the reduction of RNA structural redundancy and includes a method of selecting representative RNA structures from groups of similar structures (Leontis and Zirbel 2012). The underlying algorithm updates the database weekly and considers all RNA-containing structures deposited into the PDB. It creates integrated functional elements (IFEs), which are units of one or more RNA chains. If two RNA chains from the same structure do not interact to an appreciable extent, they are assigned separate IFEs. For example, the 28S rRNA and 18S rRNA of PDB ID 8GLP, a structure of the 80S ribosome (Holm et al. 2023), belong to two separate IFEs because they interact relatively little with one another. Meanwhile, the 28S rRNA and 5.8S rRNA of this structure are assigned to the same IFE because there is sufficient interaction between the two RNA chains. IFEs are sorted into equivalence classes, where they are grouped with other IFEs that are from the same species and exhibit similar structures. For instance, many IFEs containing the 16S rRNA of *Escherichia coli* ribosomes are grouped into the same equivalence class. A representative IFE for each equivalence class is selected by the Leontis and Zirbel method based on a calculated composite quality score that considers various metrics of the experimental structure; further details can be found at http://rna.bgsu.edu/rna3dhub/nrlist.
*(ii) Dataset used for this work.* For this study, our workflow considered the representative IFE from each equivalence class, using the 3.386 release (dated May 7, 2025) of the Representative Sets of RNA 3D Structures, referred to as the “Representative Dataset” in our paper. Several resolution thresholds were available, and we chose the 3.0 Å cutoff, which affords 1,832 high-quality RNA-containing structures and includes 2,170 representative IFEs. Notably, our workflow analyzed 253,494 amines from this dataset. While initially constructing our workflow, we worked with release 3.326 (dated March 13, 2024) which provided 197,765 amines from 1,653 high-quality RNA-containing structures split into 1,934 representative IFEs. Thus, there was a 28% expansion in the number of amines to analyze within just a little over a year’s time. When preparing a new release, the Representative Sets of RNA 3D Structures algorithm only considers a single version, often the first, of any given PDB entry. Meanwhile, our workflow uses the current version of PDB entries when it retrieves the data. Consequently, there are some changes between the structures considered by the database algorithm and those retrieved by our workflow. While it is important to mention, we do not expect that these differences in PDB entry versioning to meaningfully impact our results.
*(i) (iii) Limitations.* It is noteworthy that while the Representative Dataset has reduced redundancy, some still exists. For instance, there are 26 representative IFEs that contain the eukaryotic large subunit rRNA, with Rfam ID RF02543 (Ontiveros-Palacios et al. 2025), which have 1,000 or more observed nucleotides, according to the full CSV that was downloaded using the Representative Sets of RNA 3D Structures API. This is because each of the 26 corresponding equivalence classes is associated with a different species. Of course, even if all redundancy were removed, the dataset would still only have those RNAs whose structures have been solved, reflecting the preferences of the structural biology community.

### Computational Approach

*(i) The workflow.* We created a computational workflow that uses Python to collect and process data obtained from the structures; additionally, the R programming language was used for data analysis and plotting. Key Python libraries used in the workflow for structural analysis include PyMOL (The PyMOL Molecular Graphics System, Version 3.0 Schrödinger, LLC.), Biopython (Hamelryck and Manderick 2003; Cock et al. 2009), and GEMMI (Wojdyr 2022). The Snakemake program was used to manage our workflow. It improves reproducibility and makes computational pipelines scalable, especially when used with high performance computing (Mölder et al. 2021). The workflow can be found in its associated GitHub repository at https://github.com/The-Bevilacqua-Lab/dual-donating-amines.
*(ii) Mining and identifying dual-donating amines.* The workflow was used to iterate through each A, C, and G in the representative IFEs and identify hydrogen bonds donated by their exocyclic amine. Each structure was retrieved from the PDB, any hydrogens present were removed, and then hydrogens were added to the amines of chains specified by the IFE using PyMOL. Problematic A, C, or G residues (e.g., a G(N2) not having two hydrogen atoms) were excluded from further consideration, totaling 17 residues in this case. The code identified acceptors that reside within 4.1 Å of the amine nitrogen and that belong to polymers (e.g., RNA and proteins) but not to the parent nucleobase. The distance between each amine hydrogen and the nearby acceptor was then measured. Also, the angle between the amine nitrogen, each amine hydrogen, and the nearby acceptor was measured. Because each amine bears two hydrogens, two distance and angle measurements were made for every identified amine-acceptor pair. We expected that no more than one of these two would form an H-bond with reasonable geometry.

H-bond distance and angle criteria were determined after visualizing a heatmap depicting distance and angle measurements of the amine-acceptor pairs from the Representative Dataset (Figure 2A). The A(N6)-U(O4), C(N4)-G(O6), and G(N2)-C(O2) pairs—where N6, N4, and N2 represent the exocyclic amines—were excluded from this plot owing to their prevalence in canonical base pairing. Likewise, the putative G(N2)- C(N3) pair was also excluded from this heatmap because of its proximity to G(N2)-C(O2) H-bond that occurs in GC base pairs. Including these four pairs in the heatmap would result in tight, dense regions that hide the counts of many H-bonds that do not originate from canonical base pairing. The H-bonds from other amine-acceptor pairs exhibit a wider range of values, and we wanted to take this into account when determining the H-bond criteria. Importantly, these four amine-acceptor pairs were included in the other analyses presented in this study. After establishing the H-bonding criteria, each amine-acceptor pair was designated as H-bonding if the hydrogen-acceptor distance and the nitrogen-hydrogen-acceptor angle met the criteria.

Nucleobase amines were classified as non-, single-, or dual-donating as a function of how many H-bonds they donate. Three different scenarios can yield amines that are classified as single-donating (Supplemental Figure S9A): 1) Amines that donate one H-bond, 2) amines that donate two H-bonds where both involve the same acceptor, and 3) amines that donate two or more H-bonds where all H-bonds involve the same hydrogen. While the latter two scenarios technically involve more than one H-bond, the capacity of the amine to donate two separate H-bonds is not utilized; rather, the multiple H-bonds arise through H-bond bifurcation or trifurcation (Steiner 2002; Feldblum and Arkin 2014). We considered dual-donating amines to be those that donate two or more H-bonds where both amine hydrogens and two or more different acceptors are involved (Supplemental Figure S9B). For each dual-donating amine, the pair of acceptor atoms that participate in the H-bonding interactions were identified. In cases where a dual-donating amine had three or more acceptors, only the acceptor that resulted in the greatest H-bonding angle— which generally increases with H-bond strength—was included as the member of the amine’s acceptor pair (Steiner 2002).

1. *(iii) Canonical GC base pairs.* We occasionally considered the data in the context of GC base pairing when a G or C was involved in the dual-donating amine interaction. There are a variety of ways to infer base pairing from structural data, including the consideration of coplanarity as is performed by the DSSR software (Lu et al. 2015). In our work, GC base pairs were considered to be present when both the G(N2)-to-C(O2) and C(N4)-to-G(O6) H-bonds were formed.
2. *(iv) Visualizing instances of dual-donating amines using PyMOL.* Included in the Supplemental Material are 11 CSV files that list all identified cases related to the interactions depicted in Figure 4 and Supplemental Figure S1. Three additional CSV files are also included that list cases corresponding to Location 1 and Location 2 of Figure 7C. Supplemental Table S1 provides a brief description for each file. A Python script named “review_amines.py” is included with the workflow available from the GitHub repository. The 14 CSV files were used as inputs to review_amines.py, creating the 14 corresponding Python files that are intended to be run using PyMOL. Use of PyMOL with these Python files enables visualization of all the instances of dual-donating amines listed in the original CSV files.

Brief instructions on how to use these 14 Python files are provided here. Within the PyMOL application, click on “File”, “Run Script…”, and then the name of the Python file that you want to run. Each dual-donor-to-acceptor-pair instance (e.g., A(N6) to G(N3)/N(O2′)) is associated with an amine number that ranges from 1 to the total number of instances (e.g., 1,518). Type and enter “n” or “p” in the PyMOL command line to proceed to the next or previous dual-donor-to-acceptor-pair instance, respectfully. Type and enter “goto <NUM>” to proceed to a specific instance, where <NUM> corresponds to the amine number, which is the first column in the CSV file. We used version 3.0.0 of open-source PyMOL, but these files may work with other PyMOL versions as well.

1. *(v) Solvent accessible surface area calculations.* Biopython was used to perform solvent accessible surface area (SASA) calculations on the nitrogens of the A, C, and G amines (Hamelryck and Manderick 2003; Cock et al. 2009). For each RNA-containing structure downloaded from the PDB, the code first loads and inspects the structure for the presence of hydrogens. If any hydrogens are present, the structure is saved to a new file without the hydrogens, and then this new structure is reloaded. The SASA of each nitrogen is calculated using the method of Shrake and Rupley with the default parameters provided by Biopython, including a probe radius of 1.40 Å (Shrake and Rupley 1973).
2. *(vi) Heavy atom density calculations.* The densities of heavy (i.e., non-hydrogen) atoms within two regions of interest (ROIs), which encompass each A, C, and G amine nitrogen, were calculated (Figure 5B). First, the code worked with PyMOL to count the number of heavy atoms from polymers (e.g., RNA and proteins), as defined by PyMOL, within 7.5 and 12.5 Å of each nitrogen. The density in ROI 1 was then calculated by dividing the number of heavy atoms ≤ 7.5 Å from the amine by the volume of a sphere with a radius of 7.5 Å. For ROI 2, the two heavy atom counts were subtracted to find the number of heavy atoms > 7.5 Å and ≤ 12.5 Å from the amine. The density was then determined by dividing this number by the difference in volumes of two spheres with radii of 7.5 Å and 12.5 Å.
3. *(vii) Dihedral measurements.* The values of the *χ*, η, and θ RNA dihedrals were calculated to study nucleobase orientations relative to their attached sugars (*χ* dihedral) and sugar-phosphate backbone conformations (η and θ dihedrals). For the *χ* measurements, PyMOL was used to acquire the dihedral along the O4′, C1′, N9, and C4 atoms for purines and along the O4′, C1′, N1, and C2 atoms for pyrimidines (Bloomfield et al. 2000). The η and θ dihedrals are pseudo-torsions that involve the P and C4′ atoms of the residue in question and its immediate neighbors. These measurements were obtained using the NaTorsion program from the AMIGOS III repository (Shine et al. 2022).
4. *(viii) Calculation of p-values.* The SASA distributions of non-, single-, and dual-donating amines were compared using pairwise Wilcoxon rank sum tests, and the resulting p-values were adjusted to control the false discovery rate (FDR) (Supplemental Figure S2A) (Benjamini and Hochberg 1995). This approach was also used to compare the distributions of heavy atom densities (Figure 5C and Supplemental Figure S2C). For the distributions of χ dihedrals, which are circular, Watson’s test for homogeneity was used. Because this test generated p-value ranges (not discrete p-values), control for FDR was not performed (Supplemental Figure S3). All calculations were made using the R programming language.
5. *(ix) Clustering analysis.* The η and θ pseudo-torsions were plotted as heatmaps for non-, single-, and dual-donating amines to reveal differences between them (Figure 7A-C), and locations with higher counts in the heatmaps for dual-donating amines were identified. Some of the residues in these locations were visualized manually using PyMOL and appeared to be involved in S-motifs. Residues from each of the identified locations that correspond to a top acceptor pair category/categories (Figure 7D) were selected for further study, as specified in the Results. For each of these residues, a 6x5 nucleotide fragment was extracted from the parent structure containing the residue of interest and other nearby residues that potentially make up an S-motif. Fragment alignments and clustering analyses were then carried out to better understand the extent to which residues in the identified locations are involved in S-motifs.

For fragment alignments of each of the locations identified above, we performed RMSD calculations between all the fragments using Biopython’s Superimposer module (Hamelryck and Manderick 2003; Cock et al. 2009), and an RMSD distance matrix was constructed. To ensure equal numbers of atoms were present within each fragment, despite different sequences, a coarse-grained approach was adopted (Ramakers et al. 2024). In this approach, each nucleotide was represented using five atoms: P, C4′, N9, C2, and C6 for purines, and P, C4′, N1, C2, and C4 for pyrimidines. Hierarchical clustering analyses were then performed using the scikit-learn library (Pedregosa et al. 2011). To assess possible RMSD cut-offs for sorting fragments into clusters, we calculated three clustering validation metrics: the Silhouette score, the Calinski-Harabasz index, and the Davies-Bouldin index (Supplemental Figure S4). These metrics were calculated across a range of RMSD cut-offs determined from the linkage matrix: the smallest cut-off was the value where at least one cluster with four or more members was formed, and the largest cut-off was the integer below the maximum distance present.

The optimal RMSD cut-off was chosen from the Silhouette score because it resulted in a more manageable number of clusters. The cluster with the highest number of members was designated as the dominant cluster (Supplemental Figure S5). Within each of the resulting clusters, a representative structure was selected, which was done by finding the cluster’s average structure, calculated as the mean of all coarse-grained atomic coordinates within the cluster, and then choosing the structure with the smallest RMSD to it (Figure 7E and Supplemental Figure S6).

## Acknowledgements

The authors would like to thank Craig Zirbel for his valuable advice on the use of the Representative Sets of RNA 3D Structures database and comments on this manuscript. We also thank Bevilacqua lab members for providing helpful feedback on the manuscript, especially Dr. Elizabeth Jolley, Reuben Kern, and Kobie Kirven. Computations for this research were performed on the Pennsylvania State University’s Institute for Computational and Data Sciences’ Roar Collab supercomputer.

